# Automated video tracking and flight analysis show how bumblebees solve a pattern discrimination task using active vision

**DOI:** 10.1101/2021.03.09.434580

**Authors:** HaDi MaBouDi, Mark Roper, Marie Guiraud, James A.R. Marshall, Lars Chittka

**Affiliations:** Department of Computer Science, University of Sheffield, Sheffield, UK; Neuroscience Institute, University of Sheffield, Sheffield, UK; School of Biological and Chemical Sciences, Queen Mary University of London, London, UK; Drone Development Lab, Ben Thorns Ltd, Colchester, Essex, UK; INSECT Lab, Zoology department, Stockholm University, Stockholm, Sweden

**Keywords:** Keywords active vision, *Bombus terrestris*, cognitive strategy, flight analysis, scanning behaviour, visual recognition

## Abstract

Active vision, the ability of the visual system to actively sample and select relevant information out of a visual scene through eye and head movements, has been explored in a variety of animal species. Small-brained animals such as insects might rely even more on sequential acquisition of pattern features since there might be less parallel processing capacity in their brains than in vertebrates. To investigate how active vision strategies enable bees to solve visual tasks, here, we employed a simple visual discrimination task in which individual bees were presented with a multiplication symbol and a 45° rotated version of the same pattern (“plus sign”). High-speed videography of unrewarded tests and analysis of the bees’ flight paths shows that only a small region of the pattern is inspected before successfully accepting a target or rejecting a distractor. The bees’ scanning behaviour of the stimuli differed for plus signs and multiplication signs, but for each of these, the flight behaviour was consistent irrespective of whether the pattern was rewarding or unrewarding. Bees typically oriented themselves at ~±30° to the patterns such that only one eye had an unobscured view of stimuli. There was a significant preference for initially scanning the left side of the stimuli. Our results suggest that the bees’ movement may be an integral part of a strategy to efficiently analyse and encode their environment.

**Summary statement:** Automated video tracking and flight analysis is proposed as the next milestone in understanding mechanisms underpinning active vision and cognitive visual abilities of bees.

## Introduction

Bees are capable of memorising and discriminating a wide variety of visual patterns, including complex ones that, for example, include different stripe orientations in each of four quadrants (Benard et al., 2006; Srinivasan, 1994; Srinivasan, 2010; Stach et al., 2004; Turner, 1911; Von Frisch, 1914; Wehner, 1967). On the other hand, there is a long history of observations that bees are incapable of discriminating some relatively simple patterns (Avarguès-Weber et al., 2012; Hertz, 1929; Hertz, 1935; Horridge, 1996; Srinivasan, 1994; Von Frisch, 1914). As one example, it was reported that honeybees (*Apis mellifera*) were not able to distinguish a “plus pattern”, made up of a vertical and horizontal bar, from the same pattern rotated through 45° *i.e.* multiplication symbol (Horridge, 1996; Srinivasan, 1994) in a Y-maze setup where the patterns were displayed at a fixed distance from the bees’ decision point. Given the otherwise impressive capabilities of bees in recognising complex visual patterns (Avarguès-Weber et al., 2011; Dyer et al., 2005; Srinivasan, 1994), the difficulty in solving the plus versus multiplication sign discrimination task by bees is surprising. There is evidence that the successes and failures of bees in discriminating visual patterns are not strictly related to pattern complexity, but to the visual scanning procedures that bees use when examining and memorising the patterns (Lehrer and Srinivasan, 1994).

In vertebrates, the repertoire of such active vision strategies is already well researched (Land, 1999; Land and Nilsson, 2012; Yarbus, 2013). To scan visual targets, there can be large scale movement by the body or head, or smaller scale movements of the eyes (saccades) (Juusola et al., 2017; Najemnik and Geisler, 2005; Yang and Chiao, 2016). Such active vision is essential to obtain an accurate three dimensional representation of the material world (Kagan, 2012; Martinez-Conde and Macknik, 2008; Martinez-Conde et al., 2013; Werner et al., 2016). In some vertebrates, eye movements are also used as a sampling strategy, generating fine spatial information and improving the encoding of high spatial frequency of natural stimuli (Anderson et al., 2020; Kuang et al., 2012; Rucci and Victor, 2015). Some animals adopt a characteristic route during a visual task to facilitate target recognition (Chittka and Skorupski, 2017; Dawkins and Woodington, 2000). For instance, pigeons took stereotyped approach paths when learning to discriminate visual patterns (Dawkins and Woodington, 2000; Theunissen et al., 2017). Interestingly, they failed at these tasks when they were prevented from using their developed route. Also, characteristic head movements were observed in pigeons when stabilizing the image for forward locomotion (Theunissen and Troje, 2017).

In insects with their miniature brains, and thus possibly more limited parallel processing, there might be an even stronger need to acquire spatial information by sequential scanning than in large-brained animals (Chittka and Niven, 2009; Chittka and Skorupski, 2011; MaBouDi et al., 2020; Spaethe et al., 2006). Indeed in bumblebees, there is evidence that complex patterns cannot be discriminated when they are only briefly flashed on a screen, preventing bees from sampling in a continuous scan (Nityananda et al., 2014). Furthermore, bees exhibit defined sequences of movements in response to particular visual stimuli (Collett et al., 1993; Guiraud et al., 2018; Lehrer and Srinivasan, 1994; MaBouDi et al., 2020; Werner et al., 2016).

Here we return to one of the pattern discrimination tasks that reportedly are challenging or impossible for bees (Srinivasan, 1994) the plus versus multiplication sign discrimination task. We examine whether, and more importantly, how bumblebees can solve it. By recording the bees’ flight trajectories, and analysing their scanning movements, we aimed to determine the strategies employed in solving this visual task, specifically to investigate whether they are able to develop an active sampling strategy to solve the task if they are allowed to fly as close to the patterns as they desired.

## Materials and Methods

### Animals and Experimental Setup

Twenty bees from three colonies of bumblebees (*Bombus terrestris audax,* purchased from Agralan Ltd., Swindon, UK) were used during this study. Colonies were housed in wooden nest boxes (28 × 16 × 11 cm) connected to a wooden flight arena (60 × 60 × 40 cm) via an acrylic tunnel (25 × 3.5 × 3.5 cm). The arena was covered with a UV-transparent Plexiglas ceiling (Fig. 1*A*). Illumination was provided via high frequency fluorescent lighting (TMS 24F lamps with HF-B 236 TLD ballasts, Phillips, Netherland; fitted with Activa daylight fluorescent tubes, Osram, Germany); the flicker frequency of the lights was ~42kHz, which is well above the flicker fusion frequency for bees (Skorupski and Chittka, 2010; Srinivasan and Lehrer, 1984). The walls of the arena were covered with a Gaussian white and pink pattern (MATLAB generated); this provided good contrast between the colour of the bees and the background, required for the video analysis. Sugar water was provided at night through a mass gravity feeder and removed during the day when bees were performing experiments to ensure motivation. Pollen was provided every two days into the colonies.

**Figure 1.**
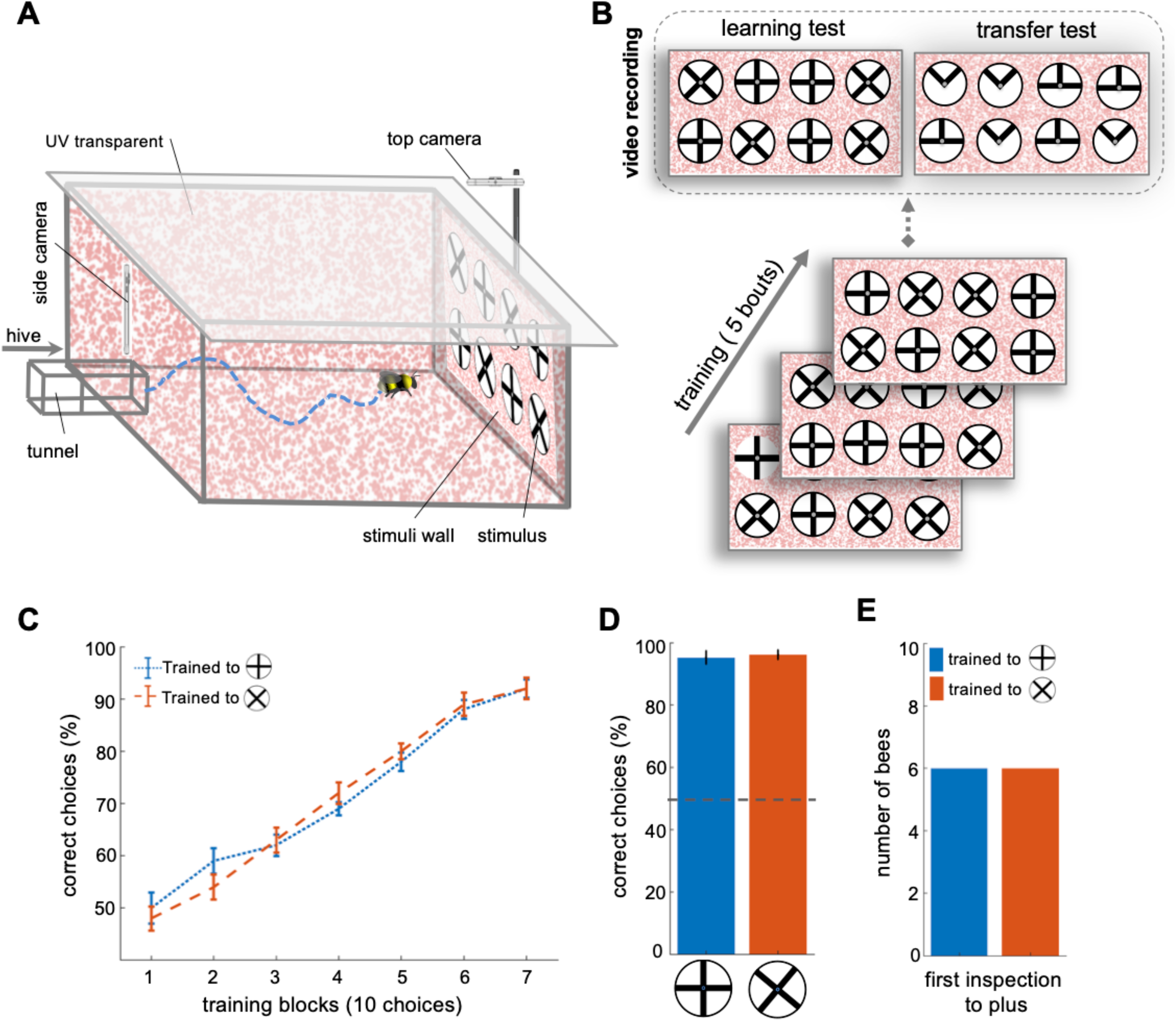
Bees’ performance in a pattern recognition task. **(A)** Schematic view of the flight arena. During the training and test experiment, an individual bumblebee was allowed to enter the flight arena through the tunnel. She then learned to fly toward the rear wall where eight patterns were displayed. She gradually learned to recognize the correct pattern (a plus or multiplication depending on the protocol) containing a sucrose solution in its centre. The flight arena was equipped with two cameras to record the bees’ flight dynamics. The side camera was placed on the top of the entrance viewing the rear wall and the top camera was placed at the top of the rear wall to record bees’ flight from above. **(B)** Training and testing protocol. The training and test patterns were constructed from 10 cm diameter white discs with 2 mm margins. Two black shapes, plus and multiplication, were presented to the bees during the training. Each pattern was attached via its centre to the rear wall of the flight arena by a microcentrifuge tube (5 mm diameter). Each bee was subjected to five training bouts, in which she entered the flight arena and was presented with eight patterns (4 plus and 4 multiplication symbols). One group of bees (n=10) was positively reinforced with the plus containing 10 μl of sucrose solution (w/w) and negatively reinforced with the multiplication containing 10 μl quinine. Another group (n=10) was trained on the reciprocal protocol. The bees were free to sample the rewarding and unrewarding patterns multiple times (refilled after departing), and return to the nest box when satiated. The place of patterns was randomly changed before each bout of the training phase. Following training, each bee’s performance was examined with two tests; here the positive and negative reinforcement were replaced with sterile distilled water. In the learning test, the bee was presented with the same patterns as during training. In the second test, the bee was confronted with the novel patterns that only displayed the upper half of the plus and multiplication symbols to the bees. One or two refreshment training bouts were used between tests to keep the bee motivation high. Bees’ flight paths were recorded for the initial 120 seconds via the two cameras. **(C)** The learning curves of two groups, blue: Group 1 (trained to plus rewarding), orange: Group 2 (trained to multiplication rewarding). Both groups of bees similarly learned to distinguish between patterns. **(D)** The performance of bees during the unrewarded learning test shows that all bees had successfully learned to distinguish between patterns (p<4.8e-3). **(E)** The number of first pattern inspections, upon entering the flight arena, that were of the plus symbol. The bees did not fly to the correct pattern from a distance (> 5 cm) more than chance (accumulated results: 10/20 correct initial visits). Blue: Group 1 (trained to plus rewarding), orange: Group 2 (trained to multiplication rewarding).

### Stimuli

The stimuli patterns were printed on laminated white discs (10 cm in diameter) to allow for cleaning (using 70% ethanol solution) in between training bouts and also tests. The training patterns consisted of two black bars (1 × 10 cm) presented in two configurations: 1) Plus pattern: one vertical and one horizontal bar aligned at their centre (⊕). 2) Multiplication pattern: same as the plus pattern but rotated by 45° (⛒). Additional patterns were constructed for a transfer test; these only presented the top half of the training stimuli (Fig. 1*B*). All patterns had 2 mm black margins around the outer circumference of the pattern. The centre of each disk was attached to the back wall of the arena via the feeder made out of a small 0.5 ml Eppendorf tube without the cap (5 mm in diameter), which contained 10 μl of either 50% sucrose solution (w/w), saturated quinine solution (0.12%), or sterilised water.

### Training and test protocol

Prior to the experiments, bumblebees could freely fly between the colony and a gravity feeder providing 30% sucrose solution (w/w) placed in the centre of the flight arena. Successful foragers were individually marked on the thorax with number labels (Opalithplättchen, Warnholz & Bienenvoigt, Germany) for identification during the experiment. Marked bees were randomly selected and pre-trained to receive 50% sucrose solution from eight white discs presented on the rear wall of the area. These pre-training stimuli were 10 cm in diameter with 2 mm wide black margins at the edges. After several bouts of pre-training, a forager that learned to take the sucrose from the feeder at the centre of the white pattern was selected for the individual experiment. During training, only the selected bee was allowed to enter the flight arena.

To improve the accuracy and the speed of learning, a differential conditioning protocol was used. Four multiplication and four plus pattern stimuli were randomly affixed to set positions on the rear wall of the arena. Each stimulus was 3-6 cm horizontally, and 5 cm vertically separated from the next stimulus, or arena wall/floor/ceiling (Fig. 1B). One group of bees (n=10) was trained to receive 10 μl 50% sucrose solution (w/w) from the feeding tubes at the centre of the plus pattern stimuli, and to avoid the multiplication patterns that contained 10 μl saturated quinine solution. The second group (n=10) was trained on the reciprocal arrangement, *i.e.* associate the multiplication pattern with a reward and avoided the plus pattern.

Bees were allowed to freely choose and feed from multiple stimuli, until they were satiated and returned to their hive; empty tubes were refilled with 10 μl of sucrose solution after the bee had left the correct stimulus and made its next choice. A bout of training was completed once the bee returned to the hive. After each bout, all feeding tubes were cleaned with soap and 70% ethanol and then rinsed with water. The patterns were separately washed with 70% ethanol. Both tubes and patterns were air-dried in the lab before reuse. The position of stimuli on the wall were randomly varied for each bout to prevent bees from using the location of the reward when solving the task.

After five bouts of training the bees were subjected to two tests, to evaluate if and how bees could recognize and select the correct pattern. In the first test, the learning test, bees were presented with the same multiplication and plus patterns used during training; this was to verify that bees had learned to associate the correct pattern with the reward, and to control for any possible olfactory cues the bees may have used during training. In the second test (transfer test), the bees were exposed to novel stimuli that only presented the top half of the multiplication and plus patterns (see Fig. 1B). This was to determine if the bees could still recognize the ‘correct’ pattern based only on the top half of the patterns. As during training, both tests provided four correct and four incorrect stimuli, randomly positioned on the rear arena wall. All stimuli feeding tubes were filled with 10 μl of sterilized water (*i.e.* no reward or punishment). One to two refreshment bouts of training (with reward and punishment) were conducted between tests to maintain the bees’ motivation. The sequence of the two tests was randomly chosen for each bee.

### Video Analysis

The arena was equipped with two cameras to record all activity of bees during tests. An iPhone 5 (Apple, Cupertino, USA) with 1280×720 pixels and 240 fps (frames per second) was positioned above the arena entrance tunnel viewing the rear stimuli wall, filming the bee’s flight in front of the stimuli wall and patterns. The second camera, a Yi sport camera (Xiaomi Inc. China) with 1280×720 pixels at 120 fps, was placed on the top of the rear wall orientated downward to view the stimuli. The first 120 seconds of each test were recorded and analysed.

To analyse bees’ scanning behaviours in front of the stimuli, prior to their choices, a MATLAB algorithm was developed that detected the bees automatically and then tracked the centroid of the bee bodies within each frame as they flew through the arena. For each frame, the algorithm subtracted a background mask image to find new candidate positions of the bee using MATLAB’s blob detection function. The parameters of this function were set to detect the blob with the same approximate size of a bee. In addition, an elliptic filter was used in the frames from the top camera to extract the bees’ body orientations. We utilised the MATLAB smoothing function (‘filter’) to exclude any erroneous data points and correct trajectories. Examples of the annotated flight paths and corresponding video recordings are shown in Figure 2 and Video S1.

**Figure 2.**
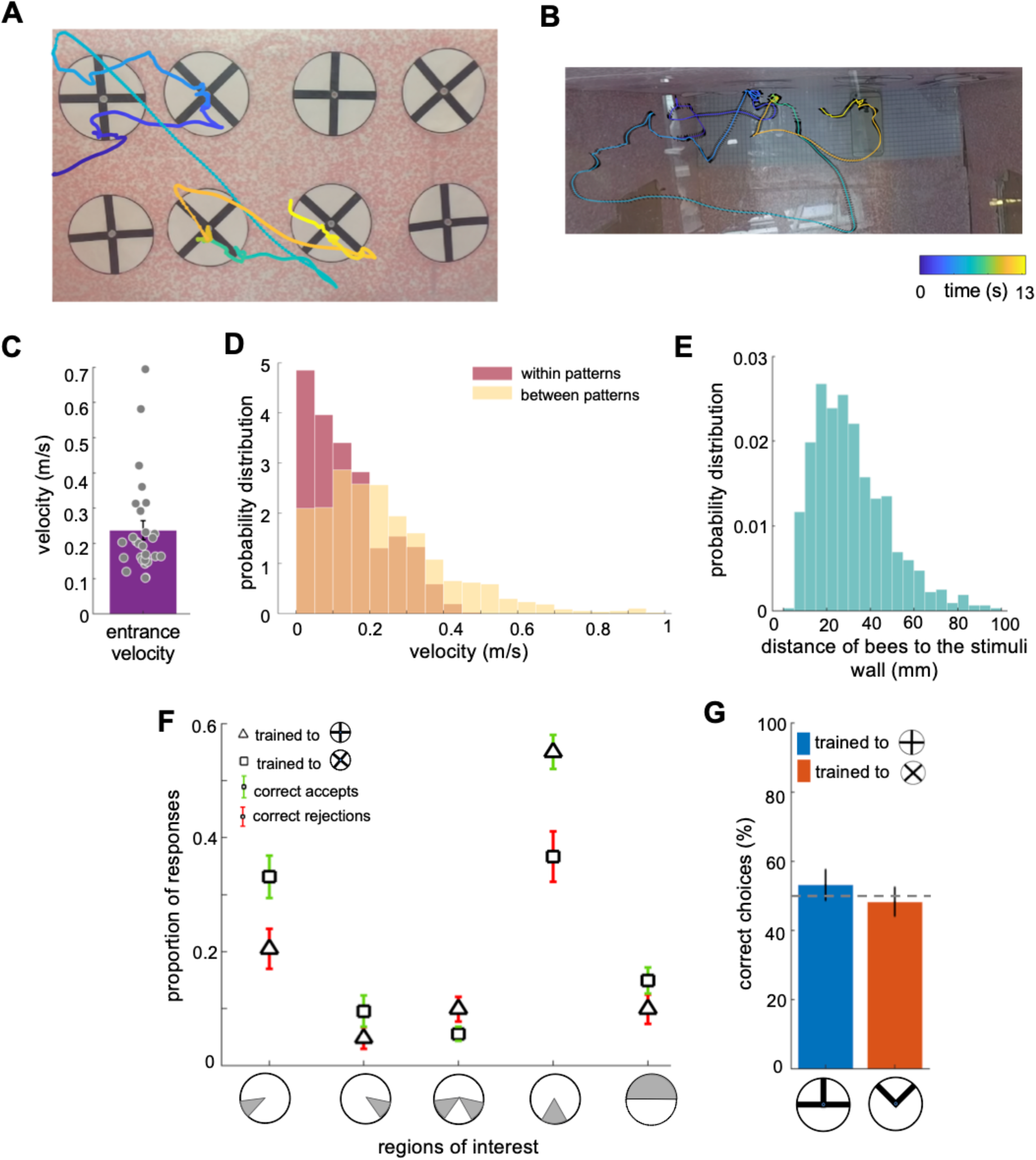
Bees’ flight analysis demonstrates an efficient strategy of scanning in bumblebees. **(A)** An example of a flight path showing the activity of a bee during part of the learning test; presented bee trained to select the plus and avoid the multiplication. Each point on the flight path corresponds to a single video frame with an interval of 4 ms between frames which was recorded from the front camera (left panel) and top camera (right panel). The bee sequentially inspected each pattern, correctly landed on multiplication and avoided the plus. The colour map changes from blue to the yellow with increasing time (See Video S1). The black lines in the left panel exhibit bee’s body yaw orientations. **(B)** Distribution of average entrance flight speed toward the wall in the learning tests (See Fig. S1). Filled dots: speed of each individual bee. **(D)** Probability distribution of the bees’ speed in two conditions; when they were inspecting patterns (Red), and when they were flying between patterns (Yellow). This indicates that they scanned patterns slower when they accepted them than when they flew to another pattern. **(E)** Probability distribution of the bees’ distance from the stimuli wall whilst inspecting patterns. **(F)** From video analysis, the proportion of scanned regions (mean ± s.e.m.) of bees’ inspections before the correct accept or correct rejection (x-axis: regions of interest are highlighted in grey). Triangles: inspection proportion (mean ±SEM) of Group 1 bees (trained to plus rewarding), squares: Group 2 bees (trained to multiplication), green error bars: correct accepts; red error bars: correct rejections. **(G)** The performance of bees during the novel test; bees equally chose both upper half-patterns (p>0.85), demonstrating that they did not learn the upper half of the patterns during training. Grey dashed lines=chance level (50%). Bar: mean performance (error bars: ± s.e.m.).

Using the first frame of each video recording, we manually specified the x, y pixel position of each of the eight pattern centres (i.e. entrances to the feeding tubes). After calculating the speed of each bee at each point of the trajectory, a threshold rule was applied to the trajectories close to the feeding tube positions to identify if the bees had landed, labelling the decision as either a correct/incorrect accept or rejection. This ‘landing’ threshold was determined by K-means clustering (MaBouDi et al., 2020a) of all bee speeds within the specified region of the feeding tubes. For further analysis of flight speeds, distances from wall, orientation, inspection times, areas of interest, and heat maps, we extracted the bees’ trajectory data (using the above procedure) from a cylindrical region in front of each stimulus, with a diameter 12 cm around the pattern centre and 2 cm out from the stimuli wall. Bespoke MATLAB algorithms were developed to calculate and plot the required datasets for each of these individual stimuli analyses (see examples: Figs. 2,3). Unfortunately one of the learning test videos from Group 1 (trained to plus) was accidently recorded at just 30 fps; we therefore excluded it from the above flight analysis. This video was sufficient, however, for the behavioural results of the choices and rejections of the bee to be extracted.

### Statistical analysis

A generalised linear mixed model (GLMM) with Binomial distribution and link function “logit” was applied to the bees’ choices recorded during the training phase to evaluate the effect of colony and group on bees’ performance and compare the learning rate between two groups of bees. To assess the bees’ performances in the tests, we analysed the proportion of correct choices for each individual bee. The proportion of correct choices was calculated by the number of correct choices divided by the bee’s total choices during the first 120s of the test. A choice was defined as when a bee touched a microcentrifuge feeding tube with her antennae or when she landed on a feeding tube. We then applied the Wilcoxon signed rank or Wilcoxon rank-sum tests to compare the bees’ responses to the learning test and the transfer test.

## Results

### Bumblebees performance in a visual recognition task

We first confirmed that bumblebees, when allowed to fly as close to the patterns as they desired, could perform the simple visual discrimination task of identifying and associating either a plus or a multiplication symbol with a reward (sucrose solution) and the other with quinine solution (penalty, bitter taste).

Bees in Group 1 recognised the plus patterns as rewarding above chance after just 20 choices (Wilcoxon signed rank test; z=2.04, n=10, p=0.04; mean=59%). Conversely, Group 2 achieved the same performance on the multiplication patterns after 30 choices (Wilcoxon signed rank test; z=2.58 n=10, p=9.7e-3). Nevertheless, there was no notable difference in the learning rate between the two groups after 30 choices (p=0.72), and the bees’ performance was not affected by colony (p=0.17) (see Table S1). The bees continued to increase in performance during the 70 choices of the training (per block of 10 choices, see Fig. 1C); whereupon all bees achieved ≥ 92% (± 7.8 s.d.) correct choice performance. The results of a generalised linear mixed model (GLMM) analysis confirmed that both groups of bees had learned to select the rewarding patterns significantly above chance (>50%) after training (Fig. 1C, p=3.84e-10). Additionally, the bees’ performance in the learning tests indicated that both groups of trained bees successfully learned to discriminate the plus from the multiplication symbol, and vice versa (Fig. 1D) (Wilcoxon signed rank test; z=2.82, n=10, p=4.8e-3 for Group 1; z=2.84, n=10, p=4.5e-3 for Group 2); again there was no significant difference between the performance of two groups in the learning test (Wilcoxon rank sum test; z=0.23, n=20, p=0.81). The bees’ performance in the leaning test was similar to that seen during the last block of 10 choices of the training phase (Wilcoxon signed rank test z=1.32, n=20, p=0.18).

During the learning tests, 6 out of the 10 bees in Group 1 (trained to plus) initially inspected (flight within 12 cm diameter of centre of pattern and 10 cm out from rear stimulus wall) the correct plus pattern. However, an equal number of bees in Group 2 also inspected the plus first, in their case the incorrect pattern. Therefore, as a whole, the bees’ pattern selection from a distance (*i.e.* from arena entrance to stimuli wall) was no different to chance (50% correct initial pattern inspections; *χ*^2^ test, Chi-square statistics*=0.8*, df=18; p=0.37) (Fig. 1E). In addition, during all the correct initial inspections, the bees still scanned the pattern before flying to the feeder tube (see Fig. S1).

### Bumblebee flight speeds and dynamics during the learning tests

To explore how bees choose the correct patterns and reject the incorrect ones, we analysed the bees’ inspection behaviours, employing a custom algorithm to track the bee locations and body orientations within each frame of the videos (Fig. 2A, B and Video S1; See Video analysis in Method section).

The bees’ initial flight speed upon entering the flight arena and approaching the first inspected stimuli was on median 0.20 (± 0.13 s.d.) ms-1 (Figs. 2C, S1). The speed reduced to a median of 0.11 (± 0.10 s.d.) ms-1 whilst in front of stimuli; the highest proportion of flight speeds was less than 0.1 ms-1 (Fig. 2D). Bees’ speed increased to a median of 0.20 (±0.24 ms-1) whilst traversing between the presented patterns. Bees typically scanned the patterns from a distance of 10 mm to 50 mm from the stimuli (Fig. 2E). The bees spent approximately 1.5 (± 0.5 s.d.) seconds in front of a stimulus, irrespective of whether this was a plus or multiplication, or the correct or incorrect pattern (Fig. 3H). The flight speed when rejecting a pattern was on average three times that of when the bee accepted a pattern and flew to the feeder. However, analysis of the flight trajectories (Fig. 3A,C,E) shows this was due to the bee accelerating away from the current pattern to the next. Interestingly, the bees showed an overall tendency to scan the patterns with their bodies oriented at ~±30° relative to the rear stimuli wall, keeping one or other eye predominantly aligned to the stimuli during the scans (Fig. 3F). Conversely, when flying between the patterns, they mostly looked forward in the direction of their motion with a much wider range of flight directions relative to the rear wall (see Discussion).

### Bumblebees scanned specific regions of the patterns prior to making a decision

As the bees did not appear to be making pattern selections from a distance (Fig. S1), we further analysed the movements of the bees whilst directly in front of the patterns. In most instances (Group 1 trained to ⊕: 89.2%, Group 2 trained to ⛒: 87%), the bees first traversed to, and then scanned, the lower part of the patterns regardless of whether the target was rewarding or aversive (Fig. 2F). Each scan led to either a landing on the feeding tube (an accept) or the bee flying to another stimulus without landing (a rejection). In Figure 2F, the proportion of bees selecting each region of the patterns prior to a decision (accept or rejection) are plotted for each group of trained bees. The highest proportion of interest was the bottom centre of the pattern with correct choices of 54.5% within Group 1, and 39.7% of correct rejections in Group 2. However, this was similar to the accumulated instances of lower left corner, lower right corner and both lower corners (summed totals for correct choices Group 2: 47.5%, correct rejections Group 1: 35%). It should be noted that the bees showed a consistent preference for the lower left corner, described further below. These preferences can be clearly seen on the heat map representation of the accumulated bee positions during scanning (Fig. 3B,D).

Bees trained on the protocol with the plus pattern as rewarding (Group 1) would typically approach the lower half of the stimulus (89% of inspections). If a plus was observed they would scan the lower centre of the pattern (containing the vertical bar) and then fly directly to the pattern centre to access the feeding tube (see Video S2). However, if the bees observed a multiplication they would usually scan the lower left corner of the pattern, containing the oriented bar of the multiplication. Of these trails, over half consisted of a single corner scan before the bees rejected the patterns. A scan of the whole pattern was clearly not required: the inspection of a single diagonal pattern element was sufficient to ascertain that the pattern was not a plus sign. In the remaining cases the bees would traverse to the opposite lower corner, then scan the remaining oriented bar before rejection (Fig. 2F). On average, only 4.5% of such inspections did the bees only scan the right corner. Bees trained on the multiplication pattern (Group 2) showed a slightly different behaviour. If the bees were inspecting a multiplication stimulus, they would first approach the left or right lower section of the pattern (see Videos S3, S4). We still observed the same preference for the left side inspections, double that of the lower right side scans. However, there were far fewer instances of bees inspecting both corners before flying to the feeding tube (Fig. 2F). When Group 2 bees (trained to the multiplication pattern) encountered a plus pattern they would again scan the lower centre at the base of the vertical bar (see Video S3). In contrast to the Group 1 bees accepting the multiplication symbol, these bees would also, on occasion, scan the lower left corner where no oriented bar was present (Fig. 3C).

**Figure 3.**
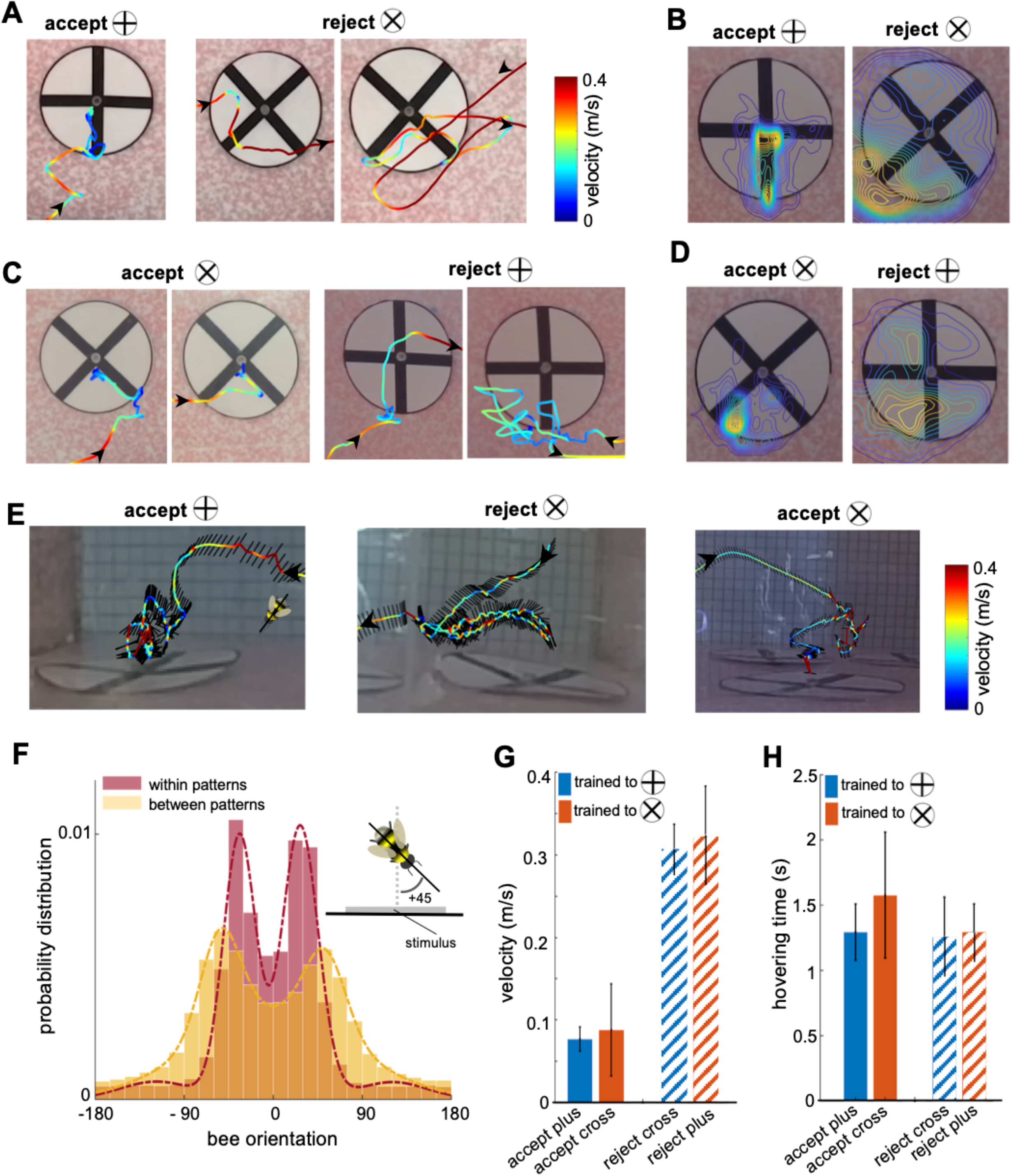
Bee scanning strategy in a pattern recognition task. **(A)** The flight paths of one example of acceptance and two examples of rejection behaviours of a bee trained to plus; the bee accepted the plus pattern after scanning the lower half of the vertical bar, while she rejected the multiplication pattern after scanning one or both diagonals bars. Line colour: flight speed 0.0 - 0.4 ms-1 (See Videos S2, S3). **(B)** Group 1 (trained to plus) probability maps (heat-maps) of bees’ locations per frame in front of plus and multiplication type stimuli during all learning tests. The yellow colours show most visited regions. **(C & D)** same analysis as A, B for Group 2 bees trained to discriminate multiplication from plus. This indicates that bees typically scanned the lower half of the pattern with a lower speed to Group 1 bees, prior to their decisions (See Videos S4, S5). **(E)** Three examples of bees’ flight paths shown from the top camera; black lines show bees’ body orientation during the flight, and arrows designate the start and ending time of scanning. **(F)** probability distribution of the bees’ body yaw orientation perpendicular to the rear stimuli wall in two conditions: when they were inspecting patterns (red) and when they were flying between patterns (yellow). Inset figure exhibits one example of bee’s orientation with +45°. This suggests the bees viewed the patterns at a median ~±30° whilst scanning, with one or other eye having a predominant view. On the other hand, when they flew to another pattern the body orientation was more parallel to the flight direction with a wider distribution of orientations relative to the stimuli wall, resulting in a median of ~±50° perpendicular to the stimuli wall. The dashed lines show the Gaussian mixture distribution models were fitted to each distribution (flights within patterns:*μ*_1_= +*27*, *μ_2_=* −*33*; flights between patterns:*μ_1_=*+*51*, *μ_2_=* −*55*). **(G)** mean flight speed (± s.e.m.) of scanning flight prior to decisions (accept and rejection) for both groups of bees. Blue: Group1 (trained to plus); orange: Group 2 (trained to multiplication). **(H)** inspection time (i.e. the time spent hovering in front of a pattern) for each symbol type for both groups of bees; inspection times of bees in front of both pattern types were equal regardless of their decision or training protocol.

To control for the possibility that the upper region of patterns may have also influenced the bees’ decisions, we carried out a transfer test (see Method section), in which bees were confronted with only the top halves of the patterns. None of the bees were able to recognize the correct pattern element, choosing equally both types of novel patterns (Fig. 2G,E) (Wilcoxon signed rank test; z=0.17, n=10, p=0.85 for Group 1; z=−0.05, n=10, p=0.90 for Group 2). We therefore concluded that bumblebees learned to only use the information of the lower sections of these patterns for recognition (similar to what is seen in honeybees (Giurfa et al., 1999), see Discussion).

These results demonstrate that bumblebees are able to learn specific features within a pattern to both accept and reject stimuli. In addition, for the specific paradigm used in this experiment, bees did not make their stimulus selection from a distance, only choosing to visit a feeder after close inspection of the presented patterns.

## Discussion

In this study, we aimed to explore the flight characteristics and active vision underpinning a simple visual recognition task in bees. Such a task was failed by honeybees (*Apis mellifera*) when they were prevented from viewing the stimuli up close (Srinivasan, 1994). Our results show that bumblebees (*Bombus terrestris audax*) can discriminate these symbols within our flight arena design. However, they chose to inspect both the rewarding and aversive stimuli from a distance of just 1 to 5 cm before making their decisions. There was no indication that the bees chose their initial stimulus from a distance when entering the arena (with 50% of initial inspections at the incorrect patterns (Fig. 1E)); even with the correctly chosen stimuli, the bees always performed a scan of the pattern elements before landing on the feeder (Fig. S1). Our experimental paradigm cannot confirm with certainty that bumblebees are unable to discriminate these simple patterns from a distance; for that we would need to control for distance as done with the honeybee experiments (Horridge, 1996; Srinivasan, 1994). However, our experiment allowed us to carefully analyse the bees’ scanning behaviour of visual features and to extract useful insights into the active vision of bees.

In brief, our bumblebees had no difficulty in learning to identify and associate either the plus or multiplication signs with reward, with all bees achieving over 90% accuracy after 70 trials (Fig. 1C). This performance was preserved during the unrewarded learning tests (Fig. 1D). Our bespoke video analysis toolkit allowed us to track the bee positions and body yaw orientations for every frame of each learning test. The most notable, and consistent, characteristics observed were:

### Partial pattern inspection

The bees primarily flew to, and scanned, the lower half of the patterns (Fig. 2F). This suggests that the lower half was all the bees learned. Indeed, when exposed to a transfer test with only the top half of the pattern available, bees failed to identify the correct halves of the training patterns (Fig. 2E). A previous study showed that honeybees (*Apis mellifera*) trained in a Y-Maze using absolute conditioning (where only the positive pattern and a secondary blank stimulus is provided) assigned more importance to the lower half of the pattern to that of the top half (Giurfa et al., 1999). During tests with only the top half of the training pattern and a novel pattern they failed to select the correct pattern half. Conversely, if bees were presented with the lower half of the training pattern and again a novel pattern they could identify the correct stimulus. In contrast, when trained using differential conditioning (using both rewarded and unrewarded patterns), the honeybees learned the whole pattern; correctly identifying both bottom and top half patterns during tests. However, in this instance, unlike in our study, the bees’ choice was recorded from a distance (for apparatus details, see (Horridge, 1996)) and bees’ flights were not analysed systematically.

In a more recent study, in which the flight path of bees was also analysed, (Guiraud et al., 2018) showed how honeybees (*Apis mellifera*) can solve a conceptual learning task of ‘above and below’ by scanning the lower of two pattern elements presented on the stimuli; this provided sufficient information for the bees to make a decision without needing to understand, or inspect, the relationship between the top and bottom pattern elements.

### Initial side preference

The bees had a significant preference for initially scanning the left side of the multiplication pattern (Fig. 2F). This left side preference for visual objects, known as pseudoneglect, is also seen in humans (Jewell and McCourt, 2000), and birds (Diekamp et al., 2005; Rugani et al., 2015). This preference may allow an individual to always start its inspection of a stimulus at the same location, allowing for consistent learning and recognition of natural stimuli; but it remains a curiosity as to why the left preference was so prevalent amongst the bees tested (Fig. 2F). In humans and birds this lateralisation of spatial attention may have evolved once in a common ancestor (Diekamp et al., 2005). However, since the visual system of insects evolved largely independently from that of vertebrates, the left-side bias must have emerged by convergent evolution. Its computational neural advantages in bees or vertebrates (if any), is not known.

### Common body orientation during scans

The yaw orientation of the bees’ bodies was most often at ~±30° to the stimuli during pattern inspections. In this manner, one or other of the bees’ eyes would face the pattern, with only a small proportion of the opposite eye having visual access to the pattern. There was no overall preference for the left or right eye (with median orientations at ~−33° and ~+27° respectively) during scans (Fig. 3F). In our previous modelling work (Roper et al., 2017), we showed that lateral connections from both the left and right lobula to the bee mushroom bodies allowed for better pattern recognition during partial occlusion of stimuli. However, this came at the expense of fine detail recognition. Therefore, and counterintuitively, having one eye mostly obscured from the pattern may provide the mushroom bodies (learning centres of the bee brain) with more distinct neural inputs. It may also allow bees to learn both the pattern and location cues simultaneously whilst scanning a resource. Future work will be needed to see if this behaviour is particular to the patterns used in this experiment, or a stereotypical behaviour.

### Commonality in scan strategies is based on stimuli, not protocol

It may seem sensible, from the bees’ perspective, if trained on plus, only to inspect the lower centre of the pattern for the vertical bar. However, both groups of trained bees initially approached and scanned the plus and multiplication in the same manner, typically checking the lower left corner of the multiplication sign and the vertical bar of the plus. This might suggest that the bees did not learn the relative position of the cues and simply searched for the first visual item at the lower left of the pattern. However, the flight tracking analysis conflicts with this hypothesis, with the inspection of the multiplication occasionally consisting of a scan of the adjacent bar of the multiplication, and with the group trained on multiplication, after scanning the vertical bar of the plus they occasionally flew to the lower left corner to presumably check for the multiplication signs oriented bar. We therefore assume that the stimulus is directing the scanning behaviour of the bee, and in turn the bee is learning both rewarding and aversive pattern features during training. Other experiments will be required to ascertain the particular rules which dictate the bees’ flight manoeuvres based on the 2D and 3D stimuli provided.

In the pioneering works of Karl Von Frisch, free-flying bees were trained to find sugar reward on certain black or coloured patterns placed horizontally on a white background (Von Frisch, 1914). Later studies showed that bees only used local cues corresponding to their approach direction when the stimuli were presented to them horizontally (Wehner, 1967). Since bees were not able to capture global shapes, this might be the reason bees could only recognise some simple patterns in the early studies. However, vertical presentation of stimuli was developed to examine what diversity of visual features bees may use, such as orientation (Van Hateren et al., 1990), radial or, bilateral symmetry (Giurfa et al., 1996; Horridge, 1996), or spatial frequency and ring-like structures (Horridge and Zhang, 1995). To control the decision distance and understand which cues were utilised by bees to recognise the target pattern, the Y-maze was introduced (Srinivasan and Lehrer, 1988). Previous research has shown that honeybees and bumblebees can solve visual tasks by extracting the localised or elemental features within the pattern (Giurfa et al., 1999; Guiraud et al., 2018). Bees may use different parts of a stimulus to discriminate between correct and incorrect patterns, depending on the training protocol employed or the specific patterns presented (Giurfa et al., 1999; Stach and Giurfa, 2005). Although the Y-maze enabled researchers to control the cues that bees could see when making decisions about visual patterns from a distance, it is a less useful paradigm to inspect the scanning strategies used by bees. Therefore, despite several decades of research in bee vision, it is still debated why, and how, bees fail to recognise some simple patterns while they show excellent recognition in other complex patterns (Avarguès-Weber et al., 2011; Dyer et al., 2005; Srinivasan, 2010). We therefore used an experimental setup in which the bees’ flight and scanning behaviour could be examined while they were close to the targets that were to be discriminated.

In this study we showcase a new suite of tools for automatic video tracking of bees in free flight and during their scanning manoeuvres, as well the algorithms needed to analyse and visualise the large amount of positional and orientation data this tracking produces. In our previous work on ‘above and below’ conceptual learning (Guiraud et al., 2018) we had to manually view and annotate 368 hours of video footage (46 hours of video footage taken at 120 fps, watched at 1/8th speed). In contrast, here the only manual process was providing a mask frame (without the bee present) per test, and marking the feeder positions within that frame. With only a small number of test videos to process this was not an issue, but even here, recent advances in making convoluted neural networks for pattern recognition accessible to non-programmers (playground.tensorflow.org, runwayml.com), as well as the more research programmer-centric DeepLabCut (Nath et al., 2019), allows researchers to provide a few dozen labelled mask frames and have these systems process thousands of mask images for all the other videos (Egnor and Branson, 2016). Similarly, the ability to visualise either individual flight paths (Fig. 3A) or combined heat maps of positional data (Fig. 3B) allowed us to quickly identify behavioural aspects of interest. Histograms of velocity, distance and orientation can be quickly generated, but more importantly the parameters defining the areas of interest can be modified and processed in a matter of minutes. Previous studies have relied upon binary fixed decision lines (Avarguès-Weber et al., 2012; Horridge, 1996; Horridge and Zhang, 1995; Srinivasan and Lehrer, 1988), with experimenters manually recording these limited behavioural data. Our in-depth analysis on such a straightforward pattern recognition task highlighted key behavioural characteristics, which can now influence future work on active vision, this simply would not have been viable without these automated tools.

## Acknowledgements

We thank Olivia Brookes for her help in collecting the preliminary data.

## Competing Interests

All authors declare no conflict of interest.

## Funding

This study was supported by the Human Frontier Science Program (HFSP) grant (RGP0022/2014), and the Engineering and Physical Sciences Research (EPSRC) Programme Grant Brains on Board (EP/P006094/1).

## Appendices

## Supplementary Figures

**Figure S1.**
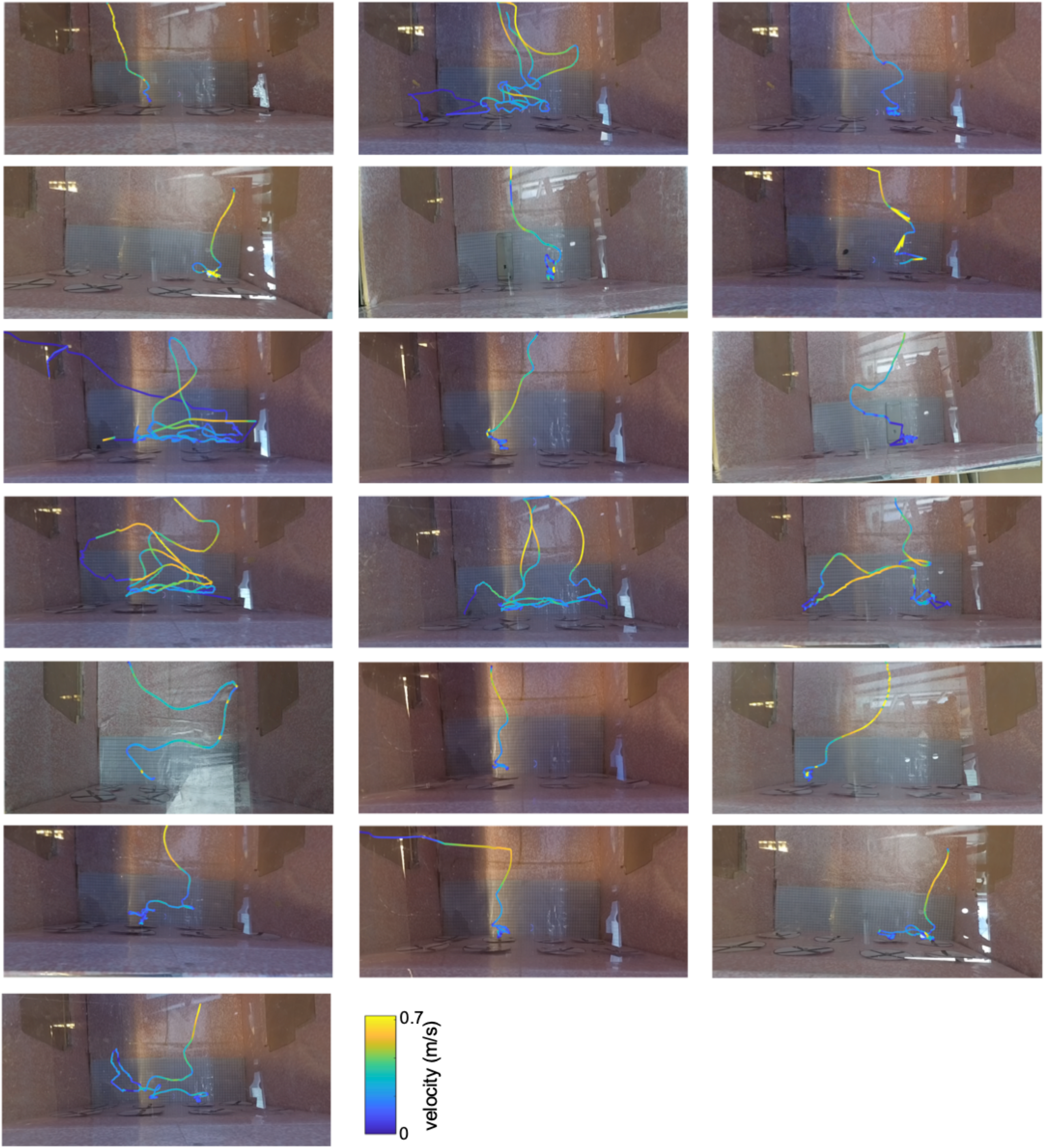
Bees’ flight paths upon first entering the arena during learning tests. In ten instances the bees initially inspected the correct stimulus, scanned the pattern and visited the feeder. In the remaining nine flights the bees initially inspected the incorrect pattern, then rejected the pattern and flew to another, usually adjacent pattern. One video is missing where the footage was only recorded at 30 fps; this bee initially inspected the incorrect pattern, and again rejected the stimulus. The bees’ first inspection appears to be random with 50/50 correct pattern selections from the arena entrance; this suggests bees have to scan the stimuli before making decisions. Line colour from blue to yellow: flight speed 0.0 − 0.7 ms^−1^.

## Notes

### Competing Interest Statement

The authors have declared no competing interest.

