## Supplementary document for "Automated video tracking and flight analysis show how bumblebees solve a pattern discrimination task using active vision"

**Running title:** ***Active scanning strategy in bumblebees***

HaDi MaBouDi[1,2,3*], Mark Roper[3,4], Marie Guiraud[3,5], James A.R. Marshall[1,2], Lars Chittka[3]

[1]Department of Computer Science, University of Sheffield, Sheffield, UK
[2]Neuroscience Institute, University of Sheffield, Sheffield, UK
[3]School of Biological and Chemical Sciences, Queen Mary University of London, London, UK
[4]Drone Development Lab, Ben Thorns Ltd, Colchester, Essex, UK
[5]INSECT Lab, Zoology department, Stockholm University, Stockholm, Sweden

***Supplementary Figures***

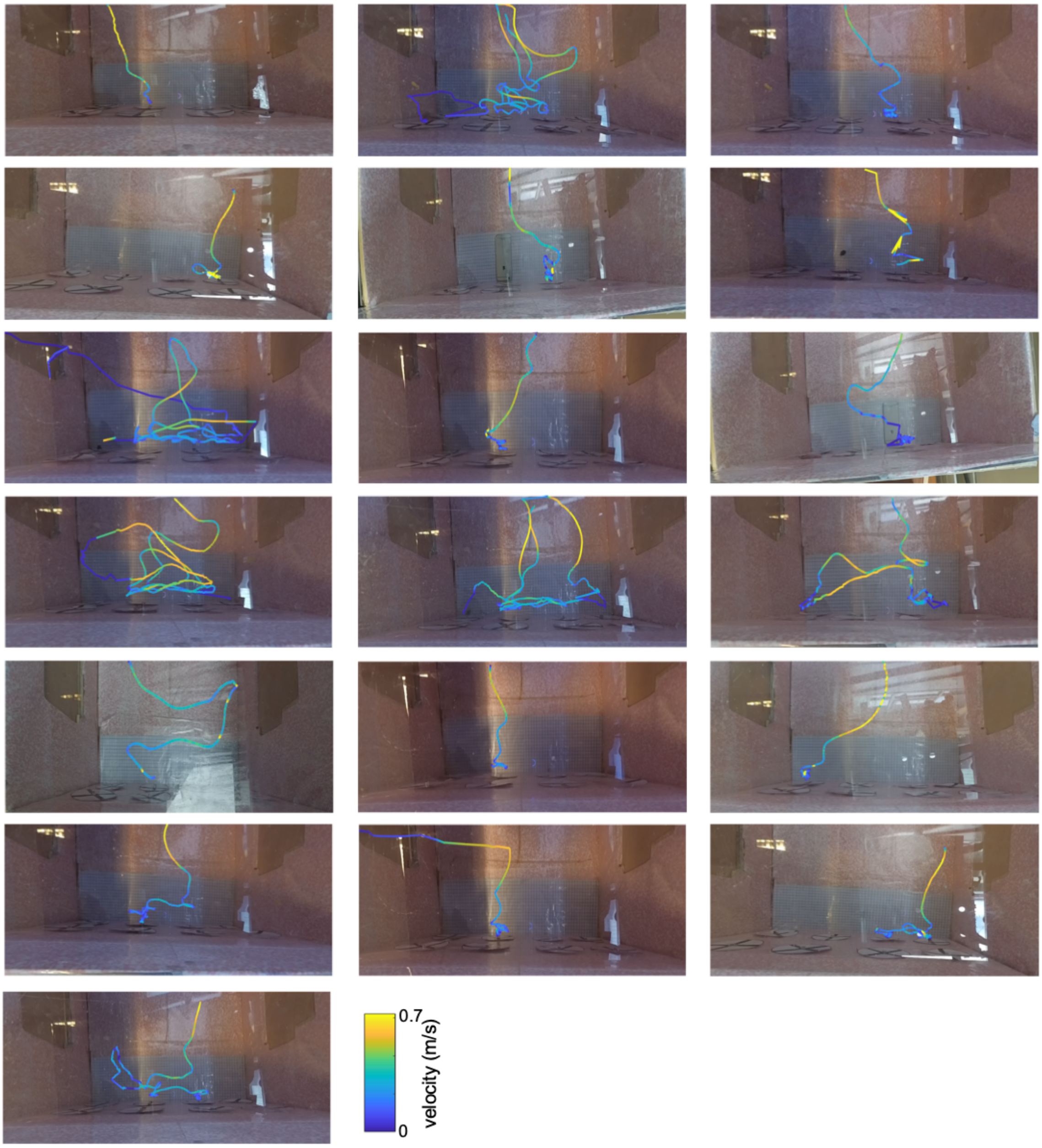


**Figure S1. Bees' flight paths upon first entering the arena during learning tests.** In ten instances the bees initially inspected the correct stimulus, scanned the pattern and visited the feeder. In the remaining nine flights the bees initially inspected the incorrect pattern, then rejected the pattern and flew to another, usually adjacent pattern. One video is missing where the footage was only recorded at 30 fps; this bee initially inspected the incorrect pattern, and again rejected the stimulus. The bees' first inspection appears to be random with 50/50 correct pattern selections from the arena entrance; this suggests bees have to scan the stimuli before making decisions. Line colour from blue to yellow: flight speed 0.0 - 0.7 $ms^{-1}$.

***Supplementary Tables:***

| Fixed factors | Estimate | SE | tStat | DF | pValue | Lower | Upper |
|---|---|---|---|---|---|---|---|
| ***Intercept*** | -0.16 | 0.50 | -0.33 | 136 | 0.74 | -1.17 | 0.83 |
| ***Colony*** | 0.21 | 0.15 | 1.36 | 136 | 0.17 | -0.09 | 0.52 |
| ***Group*** | -0.08 | 0.25 | -0.34 | 136 | 0.72 | -0.58 | 0.41 |
| ***Trials*** | 0.21 | 0.03 | 6.75 | 136 | **3.84e-10** | 0.14 | 0.27 |

**Table S1. Summary of the Generalised Linear Mixed Model (GLMM) examining factors in relation to proportion of correct choices during the training.** The dependent variable was the number of correct choices from the block of 10 choices. Fixed factors, Colony, Group and Trial were examined in the model. Bee index was calculated in the model as a random factor. Model fit statistics: AIC = 32.91; BIC=337.62; Log-Likelihood=-156.45; Deviance=312.91.

***Supplementary Videos:***

**Video S1: Example video of bee's scanning behaviour in the learning test.** The bee was trained to find the reward from the multiplication. The bee inspected lower region of the patterns and rejected the plus and accepted several multiplication signs. The video was recorded at 240 fps (frames per second).

**Video S2: Example video of rejecting the multiplication and accept the plus.** The video was recorded at 240 fps (frames per second).

**Video S3: Example video of rejecting two plus signs and accept the multiplication sign.** The video was recorded at 240 fps (frames per second).

**Video S4: Example video of from the top camera.** The video is the same as VideoS3, but recorded from the top camera. The video was recorded at 120 fps (frames per second).
